# Reward quantity discrimination in an associative learning task in wild zebrafish *(Danio rerio)*

**DOI:** 10.1101/2022.03.17.484678

**Authors:** Nawaf Abdul Majeed, Dhairyya Singh, Akshita Baiju Gopal, Tanya Battiwala, Ninaad Kulshreshtha, Rahulraj Mishra, Shagun Sabharwal, Madhusmita Behera, Manisha Sahu, Ameya Menon, Lalchhanhimi Bungsut, Amiya Walia, Raksha Saraf, Susan Mathew, Ashumi Shah, Suhaavi Kochhar, Nivedita Salar, Sushmita Thakuri, Damayanti Dasgupta, Pranaya Prakash, Arpita Ghosh, Vaishnavi Agarwal, Shubhi Pal, Preet Nagpal, Nikhil Mishra, Kiran Sahani, Rhea Lakhiani, Sahana Shanavas, Yashant Sharma, Nishtha Rampuria, Anubhab Bhattacharjee, Niharika Wagh, Sahana Hegde, Indira Bulhan, Gurasheesh Singh, Bittu Kaveri Rajaraman

**Affiliations:** Department of Biology, Trivedi School of Biosciences, Ashoka University, Sonipat, Haryana, India - 131029; Department of Psychology, Ashoka University, Sonipat, Haryana, India - 131029; Princeton Neuroscience Institute, Princeton, New Jersey, USA - 08450; University of Texas Southwestern Medical Center, Texas, USA – 75390; NISER- Bhubaneshwar, Odisha, India - 752050; Department of Psychology, University of Alberta, Edmonton, Canada -; Miranda House, Delhi, India - 110007; IISER Bhopal, Madhya Pradesh, India - 462066

**Keywords:** Reward quantity discrimination, Zebrafish, Associative learning, Quantitative cognition, Win–Stay, Lose–Shift

## Abstract

Zebrafish (*Danio rerio*) constitute an excellent model system to investigate the neural and genetic basis of quantitative cognition We tested the ability of reward quantity discrimination of wild-type zebrafish in an associative learning task. We used an operant conditioning paradigm in which the number of horizontal lines zebrafish approached in a 2-alternative forced choice task predicted the number of food reward pellets they would receive. Zebrafish did not learn a preference for the 2-line stimulus predictive of receiving 2 food pellets. However, a subset of the fish performed significantly above chance in a discrimination task with the same apparatus, in which the 2-line stimulus was associated with the same reward but the choice of the 1-line stimulus was not rewarded. Zebrafish also showed a population level learning in a simpler spatial association task with the same apparatus. We also explored the explanatory value of alternative spatial learning hypotheses such as a Win-Stay, Lose-Shift (WSLS) strategy at the individual level for fish in navigating these spatially randomised tasks.

## Introduction

Early mathematical pedagogy focuses on symbolic representation of the numerosity of objects (Dehaene and Stanislas, 1997). Our understanding of the neuronal basis of human quantitative cognition is limited due to the relatively low resolution of human imaging, and animal model systems are necessary to take forward a circuit-level understanding of quantitative cognition.

However, the skill most widely tested in non-human animals is non-symbolic quantitative discrimination, through tasks involving spontaneous or learned discrimination between sets of objects of varying cardinal numerosity. Associative reward-based learning tasks that reward the choice of stimuli of a particular cardinality have been used successfully to elicit responses in a range of animals: chimpanzees (Beran et al., 2013; Beran, Rumbaugh, et al., 1998; Boysen & Berntson, 1989; Garland et al., 2014; Matsuzawa, 1985; Murofushi, 1997; Rumbaugh et al., 1987, 1988; Tomonaga, 2008; Tomonaga & Matsuzawa, 2002; Woodruff & Premack, 1981), rhesus macaques (Beran et al., 2008; Cantlon & Brannon, 2007; Jordan et al., 2008; Livingstone et al., 2014), dogs (Macpherson & Roberts, 2013; Miletto Petrazzini & Wynne, 2016; Range et al., 2014; Ward & Smuts, 2007; West & Young, 2002), dolphins (Jaakkola et al., 2005; Kilian et al., 2003), racoons (Davis, 1984), rats (Church & Meck, 1984; Davis et al., 1989; Davis & Albert, 1986; Davis & Bradford, 1987; Suzuki & Kobayashi, 2000), mice (Berkay et al., 2016; Çavdaroğlu & Balcı, 2016; Light et al., 2019), horses (Uller & Lewis, 2009), bears (Vonk & Beran, 2012), grey parrots (Aïn et al., 2009; Pepperberg, 1987, 2006; Pepperberg & Gordon, 2005), crows (Kirschhock et al., 2021), pigeons (Emmerton et al., 1997; Emmerton & Renner, 2006; Machado & Keen, 2002; Machado & Rodrigues, 2007; Roberts & Mitchell, 1994; Scarf et al., 2011; Tan et al., 2007; Tan & Grace, 2010), robins (Garland et al., 2012; Hunt et al., 2008), canaries (Pastore, 1961), and chicks (Rugani et al., 2007, 2008, 2009, 2013, 2014). Work on other vertebrate systems such as reptiles: tortoises (Lin et al., 2021) and lizards (Petrazzini et al., 2017) and amphibians: frogs (Stancher et al., 2015), salamanders (Krusche et al., 2010) has been scarce, but there has been a mushrooming of work on associative learning of quantities in fish: zebrafish (Bisazza & Santacà, 2022; Potrich et al., 2015) and the related *Danionella* (Zanon et al., 2025), angelfish (Gómez-Laplaza & Gerlai, 2011a, 2015; Miletto Petrazzini & Brennan, 2020), archerfish (Potrich et al., 2022; Karoubi et al., 2017), goldfish (DeLong et al., 2017), redtail splitfins (Stancher et al., 2013), mosquitofish (Agrillo et al., 2009), guppies (Agrillo et al., 2014; Agrillo, Piffer, et al., 2012; Piffer et al., 2012), as well as some comparative work (Agrillo, Petrazzini, et al., 2012; Agrillo & Bisazza, 2017; Xiong et al., 2018).

Symbolic numerical representation has also been proposed as an aspect of numerical cognition that may enable precise numerical encoding even for larger numerosities in humans, and this has only been studied in a few other species such as chimpanzees (Beran, Savage-Rumbaugh, et al., 1998; Biro & Matsuzawa, 2001; Boysen et al., 1996; Boysen & Berntson, 1989; Matsuzawa, 1985), monkeys (Judge et al., 2005), parrots (Pepperberg, 2006), pigeons (Xia et al., 2001), archerfish (Karoubi et al., 2017) and bees (Howard et al., 2019b), besides humans (Feigenson et al., 2004), in comparison to the large body of work on non-symbolic numerosity.

Spontaneous quantity preference tasks have been insightful about the ecological value of general quantitative cognition and magnitude estimation. Quantitative cognition can be useful in gauging food quantity during foraging in chimpanzees (Beran et al., 2008), mice (Panteleeva et al., 2013), cats(Chacha et al., 2020), robins (Hunt et al., 2008), crows (Bogale & Sugita, 2014), guppies (Lucon-Xiccato et al., 2015), spiders (Rodríguez et al., 2015) and carnivores more generally (Benson-Amram et al., 2018); in avoiding large groups of predators in redshanks (Cresswell, 1994), minnows (Hager & Helfman, 1991), piranhas (Queiroz & Magurran, 2005), guppies (Botham et al., 2005; Lucon-Xiccato et al., 2015; Magurran & Seghers, 1994), killifish (Morgan & Godin, 1985) or even avoiding persistent harassment from sexually interested conspecific males in mosquitofish (Agrillo, Dadda, Serena, et al., 2008; Agrillo et al., 2007); in assessing group size during social or sexual aggregation in monkeys (Jordan et al., 2005), guppies (Lindström & Ranta, 1993; Lucon-Xiccato & Dadda, 2017), other poeciliids (Agrillo & Dadda, 2007), mosquitofish (Agrillo, Dadda, & Serena, 2008; Agrillo, Dadda, Serena, et al., 2008; Agrillo et al., 2007), angelfish (Gómez-Laplaza & Gerlai, 2011b), swordtails (Buckingham et al., 2007), sticklebacks (Mehlis et al., 2015; Tegeder & Krause, 1995), perch (Binoy & Thomas, 2004), zebrafish (Potrich et al., 2015; Pritchard et al., 2001; Seguin & Gerlai, 2017), and mealworm beetles (Carazo et al., 2012); or during conflict in chimpanzees (Wilson et al., 2002), lions (McComb et al., 1994), dogs (Bonanni et al., 2011); or even in contexts such as parental care in cichlids (Forsatkar et al., 2016) and searching for shelter in snails (Bisazza & Gatto, 2021). Most of these tasks deal with the animal’s ability to discriminate between the cardinality of two or more quantitatively specific stimulus sets, but ordinal cognition can also be useful for navigation. Experiments manipulating the ordinality of landmarks passed relative to distance passed suggest that ordinal information may be used instead of spatial distance by either a small fraction of individuals in some systems like honeybees (Chittka & Geiger, 1995), or by a majority of individuals for example in chicks (Rugani et al., 2007, 2008) and guppies (Petrazzini et al., 2015), or along with spatial information in zebrafish (Potrich et al., 2019).

The zebrafish model system lends itself to large-scale brain imaging with single-cell resolution (Bianco et al., 2011). Along with its strong genetic manipulation toolkit (Levin & Cerutti, 2009), this model system enables a powerful investigation of the neural circuitry underlying early quantitative cognition. Initial molecular assays of c-fos and egr-1 expression (Messina et al., 2020, 2022) and preliminary results from calcium imaging (Luu et al., 2024) have already narrowed down parts of the zebrafish brain that might be involved in quantitative cognition. This study aims to examine the ability of wild zebrafish to associatively learn non-abstract symbols representing reward quantity, building upon experiments demonstrating the use of zebrafish as a model system with non-symbolic cardinal (Agrillo, Petrazzini, et al., 2012; Bisazza & Santacà, 2022), ordinal (Potrich et al., 2019) as well as proto arithmetic abilities (Potrich et al., 2024). Our study specifically examines the associative learning of stimuli that act as predictors for the number of food pellets received as reward, and we conduct three experiments to arrive at an understanding of performance on various degrees of task difficulty.

## Material and methods

### Subjects

Experimental subjects were wild-caught zebrafish (*Danio rerio*) bought commercially and in turn sourced from local ponds and rivers in India. Details are in the Animal Welfare Note. The first four batches as well as the last batch of fish were used for Experiment 2; the second set of 11 zebrafish was used for Experiment 1. All of these fish were experimentally naive.

The first batches were housed in a generic cuboid glass aquarium (18×12×9 inches) and filled with 20 litres of RO-purified water, for at least 7 days before the experimental period began. The water in the tank was filtered through a SOBO® WP-950F 6-Watt sponge filter and the tank was maintained at 27 °C (± 1 °C) using an RSElectrical® RS008-A 50-Watt submersible water heater. The second and third batches of fish were housed in the ZebTec Active Blue - Stand Alone system (Tecniplast, PA, USA), which was auto-maintained with a pH of 7.50-8.50, 28-30℃ temperature and 650-700 µS conductivity. In all housing conditions, the fish were maintained in a 14:10 light: dark photoperiod by switching on and off a Roxin® RX-300 6-Watt fluorescent LED providing ambient light using a Smaerteefi® programmable power-strip, and later using a custom-built switch. The subjects were maintained on a daily feed of Tetra® Tertabits Complete flake food, except during the training phase when they did not receive food outside the training paradigm. The ARRIVE international guidelines and The Guide to Ethical Information Required for Animal Behaviour Papers were followed in the reporting in this manuscript. ASAB/ ABS Guidelines for the Use of Animals in Research and CSPCA national and institutional guidelines were followed for the care and use of animals. Ethical approval was obtained from our University’s Institutional Animal Ethics Committee (approval no. in Animal Welfare Note).

### Apparatus

For training and testing, a transparent glass T-Maze was purpose-built for the experiment (Figure 1). The dimensions of the stem were 20×4×15 inches (LxWxH) while the dimensions of the arms in total were 20×4×15 inches (LxWxH), with each arm’s dimensions being 8×4×15 inches (LxWxH). A sliding gate was fixed 5 inches up the stem made of transparent polycarbonate material attached to the tank with epoxy-resin adhesive (Araldite®). For the stimulus presentation, a BenQ® Zowie XL2411 24-inch 144 Hz 1080p monitor was located right in front of the maze arms and connected via HDMI to a Lenovo® S400z All-in-one PC. To record experimental trials for later analysis, a Nikon® Coolpix A900 was mounted via a Manfrotto® 290 dual tripod over the T-maze. The maze contained 5 litres of RO-filtered water, with the water level maintained just above the position of the stimuli on the screen. To provide a controlled visual environment and block the visibility of the experimenters, 24×36 inch white chart paper was used to cover the maze setup.

**Figure 1:**
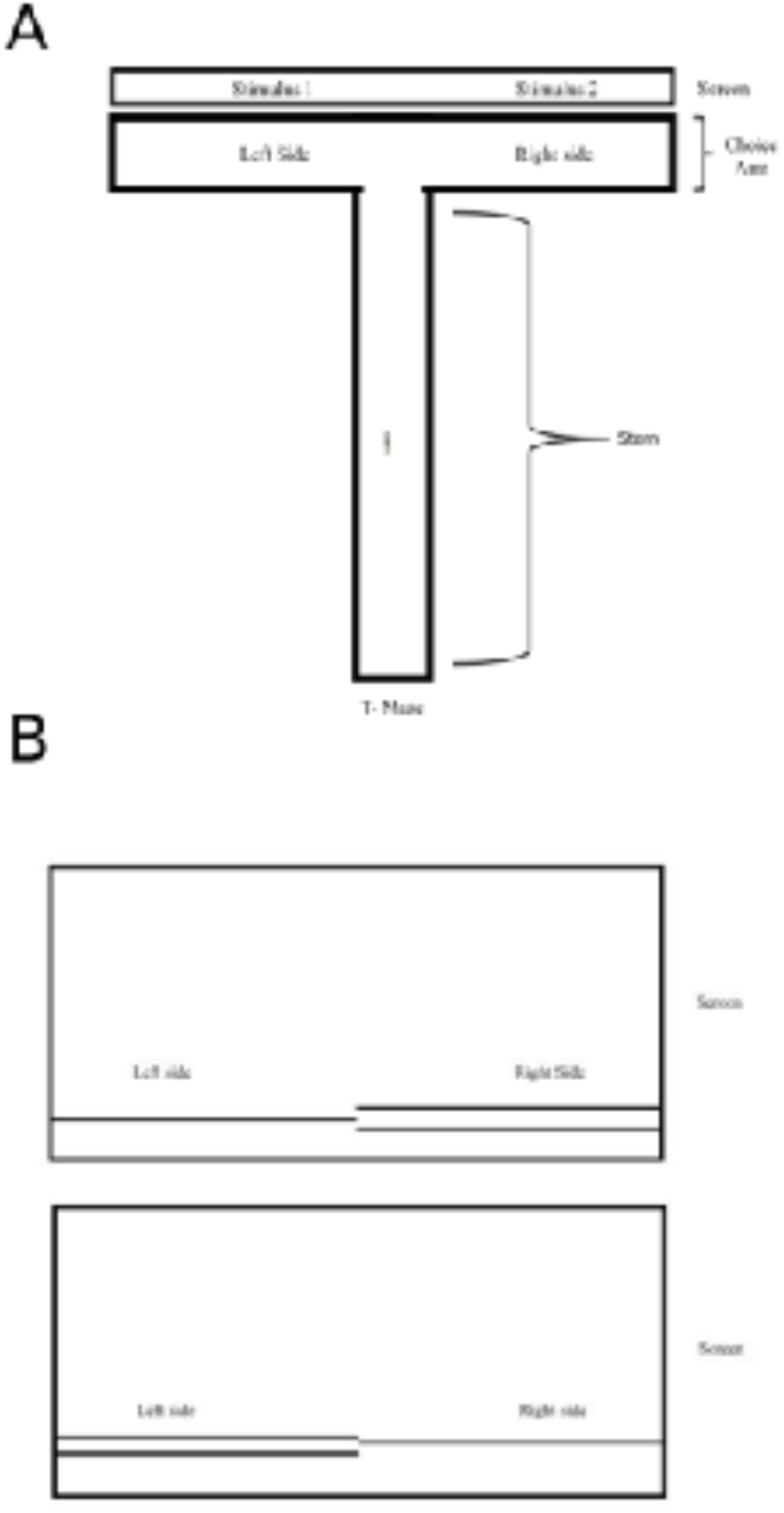
Schematic representation of the experimental set-up and stimuli. A) Schematic diagram of the T-maze with the screen attached to it (view from above). B) Visual stimuli as shown on the computer screen (as viewed by fish swimming up the trunk of the T maze).

### Stimuli

The stimuli were produced by a custom-written MathWorks® MATLAB script using Psychtoolbox. The stimuli consisted of black (RBG 000) 6-pixel thick horizontal lines drawn on an all-white background (RGB 256 256 256) in the bottom third of the screen. The side where the stimuli with two lines would be displayed was randomised between the left and right sides of the screen.

### Experiment 1: Testing protocol: associating symbols with food reward pellet quantity

The experimental protocol was divided into 3 phases – the acclimatization phase (3 days), the training phase (20 days) and the testing phase (the last and 21st day). Each day of training involved a sequence of 10 rewarded trials for the first three batches of fish. Since the performance of fish was slightly better in the first 5 trials, we experimented with a smaller number of 5 trials per day for the last batch to see if it improved performance. In both cases, the final testing day involved the same number of unrewarded probe trials as during training.

For acclimatization, the fish were transferred to the T maze using a small cup to scoop up the fish such that the fish stayed within the water with no periods of asphyxiation. Fish of both sexes were used in experiments. Our pilot trials showed initial fear responses in zebrafish when newly introduced to the T maze, either freezing or frantic swimming, and we adapted the acclimatization protocol in Sison and Gerlai (2010) to habituate the fish to the maze in decreasing-size shoals over three days before the first training session. On day 1, 4 fish were transferred together to the T-maze, sample stimuli were put up on the screen and 10 Tetra® Tertabits Complete food flakes were dropped at the ends of each T-maze arm at both ends of the left and right arm. The fish were allowed 2 hours to acclimatize together and get familiar with both the structure of the maze and the location of food. This was repeated with pairs of fish on day 2, and then for individual fish on the third day.

For training and testing sessions, individual fish were transferred into the stem of the T-maze, held in by the gate, for a 20-minute acclimatization period, after which the MATLAB stimulus script was triggered, displaying the stimuli consisting of 1 and 2 horizontal lines located randomly on either side of the screen. Then, the gate was lifted and after the fish made the first choice towards one arm of the maze, the fish was rewarded with either one food flake or two, depending on the stimulus displayed in the arm it chose. As soon as the fish consumed the reward, the stimuli were replaced with a blank white screen. For the second and third batch of fish, additional gates were closed after the fish made the first choice to ensure the reward was eaten - this made no difference to the results. For all fish, once they returned to the stem of the T maze, the gate was put back in to hold them in the distal part of the stem and effectively reposition them for the next trial. All sessions were videotaped and analyzed by a second observer who was blind to the side-stimulus association. Control trials with no displayed stimulus during acclimatization and training were also used to assess any side bias in the fish. Freezing or side-biased behaviour were used as exclusion criteria for continued training and testing.

### Experiment 2: Testing protocol: Associating symbols with a food reward

We implemented another version of our experimental protocol as a control, for a simpler task that has been studied and reported in the literature. This tested associative learning of the same visual stimuli, but instead of the number of stimulus lines predicting the number of food flakes received as a reward, there was a 2-pellet reward for the larger stimulus and no reward for the smaller stimulus. The protocol was otherwise identical to the one above.

For the task of associating visual stimuli with food reward, a sample size of N=7 was calculated as the number of individual fish required to obtain a significant effect similar to literature which conducted a similar task with zebrafish (Potrich et al., 2015, 2019). This number was obtained by conducting an a priori power analysis using the program G*power 3.1.9.4 (Faul et al., 2007) using the average effect size observed in literature for similar tasks in zebrafish (Potrich et al., 2015, 2019). The power analysis was conducted for the two-tailed Wilcoxon signed-rank test with power (1 - β) set at 0.80, effect size (d) = 1.489 and alpha = 0.05.

### Experiment 3: Testing protocol: Preference for reward quantity in a spatial association task

We performed an additional control experiment to assess food pellet quantity preferences to tackle the most simple explanation for the difference in outcomes between Experiments 1 and 2: that zebrafish are not strongly enough motivated by the difference between 2 and 1 pellets of food - the key difference between the testing paradigms used in Experiments 1 and 2. We tested this by checking whether zebrafish showed a spatial preference between these quantities of food reward, where going to one side was always rewarded with 2 food pellets and the other side was not rewarded with food.

### Statistical Methods

All statistical tests were conducted in R v.1.2.2. A Shapiro-Wilk test was conducted to check if our data were derived from a normal distribution. We conducted a Kruskal-Wallis test to check for any differences in the proportion of fish choosing the side with 2 lines across days of training in the 1 vs 2 food reward paradigm. To check for any consistent change in the proportion of trials in which the fish chose the side with 2 lines, and to pool across days to reduce inter-day stochasticity, we conducted a Kruskal-Wallis test across non-overlapping sets of 5 consecutive days. We conducted pairwise Wilcoxon tests with Bonferroni correction to test the significance of the differences in the proportion of fish choosing the side with 2 lines between groups of 5 consecutive days. We used a one sample Wilcoxon signed rank test to check for the significance of the performance of the fish in the last set of 5 consecutive days above the chance level (0.5). We used a binomial test to check the minimum required number of successful events per number of trials to define statistically significant learning criteria for individual fish.

## Results

### Experiment 1: Associative learning of quantitative symbols predictive of food reward quantity

We trained zebrafish in a 2 alternative forced choice task to associate visual stimuli consisting of 1 or 2 lines with the same number of food pellets delivered as a reward. Aggregating across all fish that completed the task (n = 24), we found no statistically significant learning of a first-choice preference for the 2-line stimulus in this quantitative association task at the whole population level. A Shapiro-Wilk test conducted on the distribution of performance across fish for each day did not uphold the approximation of the proportion of choices by a normal distribution. The median proportion of trials per day across fish in which they chose the 2-line stimulus associated with two pellets of food was 0.5 +/− 0.2 (median +/− interquartile range) on day 1 and 0.4 +\− 0.35 on day 21 (median +/− interquartile range), and it varied between 0.4 and 0.6 during the 21 days (Figure 2A). A Kruskal-Wallis test shows that the proportions of daily trials in which fish choose the side with 2 lines (indicating two pellets of food) are not the same across days for all the fish combined (p-value = 0.04703), due to the high variation in the performances of the fish across individual days. But when comparing the performance using pairwise Wilcoxon test results for a range of bins across groups of days, we found no significant learning at the population level in this task. The cumulative proportion of choices of the 2-line stimulus (Figure 2B) did not differ from the performance expected by chance alone (a probability of 0.5 in choosing either stimulus).

**Figure 2:**
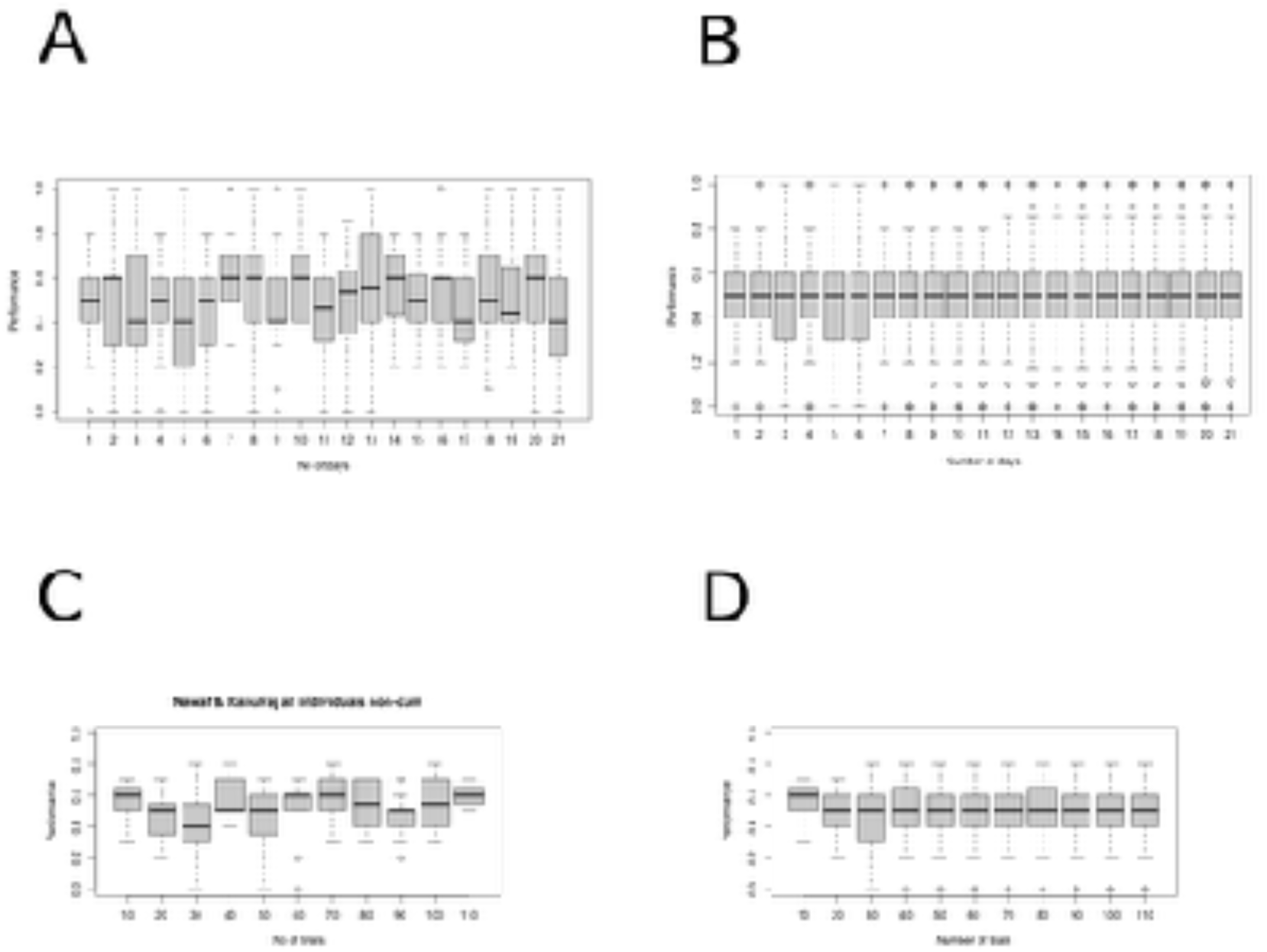
The (A, C) daily performance and the (B, D) cumulative proportion of trials in which zebrafish chose to move to the side with the 2-line visual stimulus and receive 2 pellets of food as reward, across 21 days, for all fish (n=24) (A, B), and (C, D) for the subset of fish who were individually tracked (n=11).

Initial trials did not distinguish between individual fish, although we observed considerable interindividual differences. In two batches of fish tested (n=8+3), the individual identities of fish were kept separate throughout trials, and inter-individual differences were noted. The daily (Figure 2C) and cumulative performance across trials (Figure 2D) for this subpopulation of fish also did not differ from chance expectation (0.5), as with the wider population. But for all individually tracked fish, we investigated their individual ability to reach the learning criteria used in the literature - of reaching a significant departure from chance over several consecutive trials in a binomial test. Only two out of eleven fish, R3 (Figure 3A) and M4 (Figure 3B) reached a performance of 66% correct choices over 30 trials that were significantly above chance (p-value = 0.04937), although their individual trajectories showed fluctuation about the chance line.

**Figure 3:**
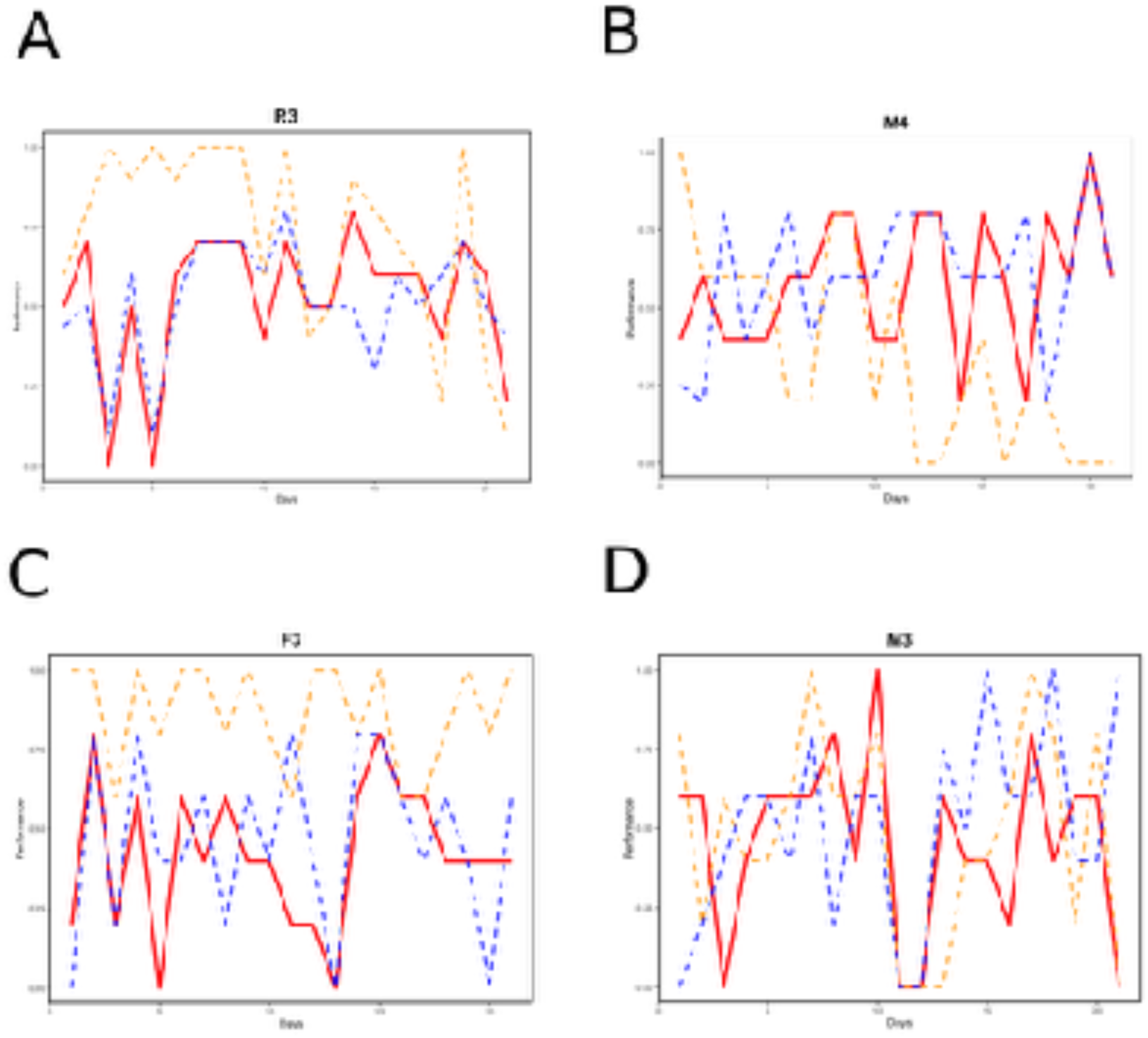
The proportion of trials for individual fish with better (A, B) and worse (C, D) performance in choosing the side with 2 lines in the 1 vs 2 food reward paradigm (thick red continuous line), plotted alongside the proportion of trials in which they chose just the right side (orange dotted lines), and the average Win-Stay/Lose-Shift scoring metric (blue dotted lines) across days of training.

We also explored alternative hypotheses to explain individual behaviour. One alternative hypothesis is that fish are side-biased rather than showing associative learning. If fish were to choose a single side of the maze to move to consistently, with the side-randomized presentation of stimulus we provided they would end up making the more rewarded choice concerning the visual stimuli 50% of the time. We plotted each individual fish’s performance relative to the magnitude of the stimulus (which in turn predicts reward magnitude) (thick red continuous line, Figure 3), alongside the fish’s proportion of choices towards the right side of the T maze (orange dotted lines), across days of training (Figure 3). In parallel, we explored another hypothesis: that fish were utilizing a Win-Stay/Lose-Shift (WSLS) strategy based on spatial learning from past rewarded trials (Byrne et al., 2020; Nowak & Sigmund, 1993; Rayburn-Reeves et al., 2013). Since the fish encounters a fluctuating reward relative to any given side, this hypothesis posits that fish continue to choose a particular side if it is rewarded in the previous trial, but would switch sides in the absence of reward. We calculated an extension of this WSLS scoring metric (plotted in blue dotted lines, Figure 3) where the fish’s behaviour is given a score of 1 if the fish returns to the last visited side and received the higher reward with 2 pellets of food, OR if it avoids the last visited under-rewarded side with 1 pellet. The fish gets a score of 0 if it visits the same side as before despite getting a lower reward in the previous trial, or if it avoids the last visited side even if that led to a higher reward.

We found inter-individual differences not only in performance on the task but in side bias and the Win-Stay/Lose-Shift score of their choices. Of the two fish which performed significantly above chance levels, fish R3 seemed to initially have a right-side bias which reduced later on (Figure 3A); while fish M4’s rise in the proportion of trials choosing the 2-line stimulus over time can be explained by a Win-Stay/Lose-Shift strategy (Figure 3B). Many other fish that were individually tracked were side biased throughout, such as F3, which used a right-side biased approach relatively consistently after the first two days of the experiment, with a relatively low score in learning the association between stimulus and reward quantity (Figure 3C). A combination of the side bias and WSLS score explains the fish M3’s choices from trial to trial better than the fish learning an association between the stimulus and the reward quantity it represents (Figure 3D).

### Experiment 2: Associating quantitative symbols with the presence/absence of a food reward

Given that these zebrafish showed no population level learning of this associative learning task between quantitative symbols and reward quantity, and that our protocol differed on several counts from the protocols used in the literature to assess quantitative cognition: utilizing locally wild-caught zebrafish, a T-maze and computerised horizontal line stimuli, we attempted to replicate a simpler task documented in the zebrafish literature (Agrillo, Petrazzini, et al., 2012; Bisazza & Santacà, 2022) to enable us to assess the performance of zebrafish on a documented task in our setup.

Using the same set-up, we assessed the performance of zebrafish in learning to choose the side of the tank displaying the 2-line stimulus, with a simpler reward structure: the 2-line stimulus was rewarded with 2 pieces of food, and the 1-line stimulus with an absence of food reward (2 vs 0 reward), rather than with the number of lines being associated with the quantity of the food reward (2 vs 1 reward).

Two of these individually tracked fish showed freezing behaviour, and the remaining population of 10 fish did not show a stable or statistically significant departure from chance (Figure 4A, 4B). Many more individuals showed freezing behaviour as trials progressed, or statistically significant side bias, leading to their exclusion from the experiment within the first few days of training. A subpopulation of five fish showed a reduction in side bias and continued to learn with a gradual but unstable elevation in the day-by-day performance, rising further along with a reduction in performance variability in the last few days (days 26-30) (Figure 4C). The proportion of trials in which these fish chose the 2-line stimulus associated with the presence of reward was 0.4 +/− 0.2 (median +/− interquartile range) on day 1 and 0.8 +\− 0.00 on day 30 (median +/− interquartile range). The cumulative performance of the fish across all trials stabilized around day 14 at 0.6 when looking at the cumulative proportion across all trials (Figure 4 D). A one-sample Wilcoxon signed-rank test also showed that the proportion of choosing the side with the 2-line stimulus for these days 26-30 is significantly higher than the chance level (0.5) for this group (p-value = 0.0002406).

**Figure 4:**
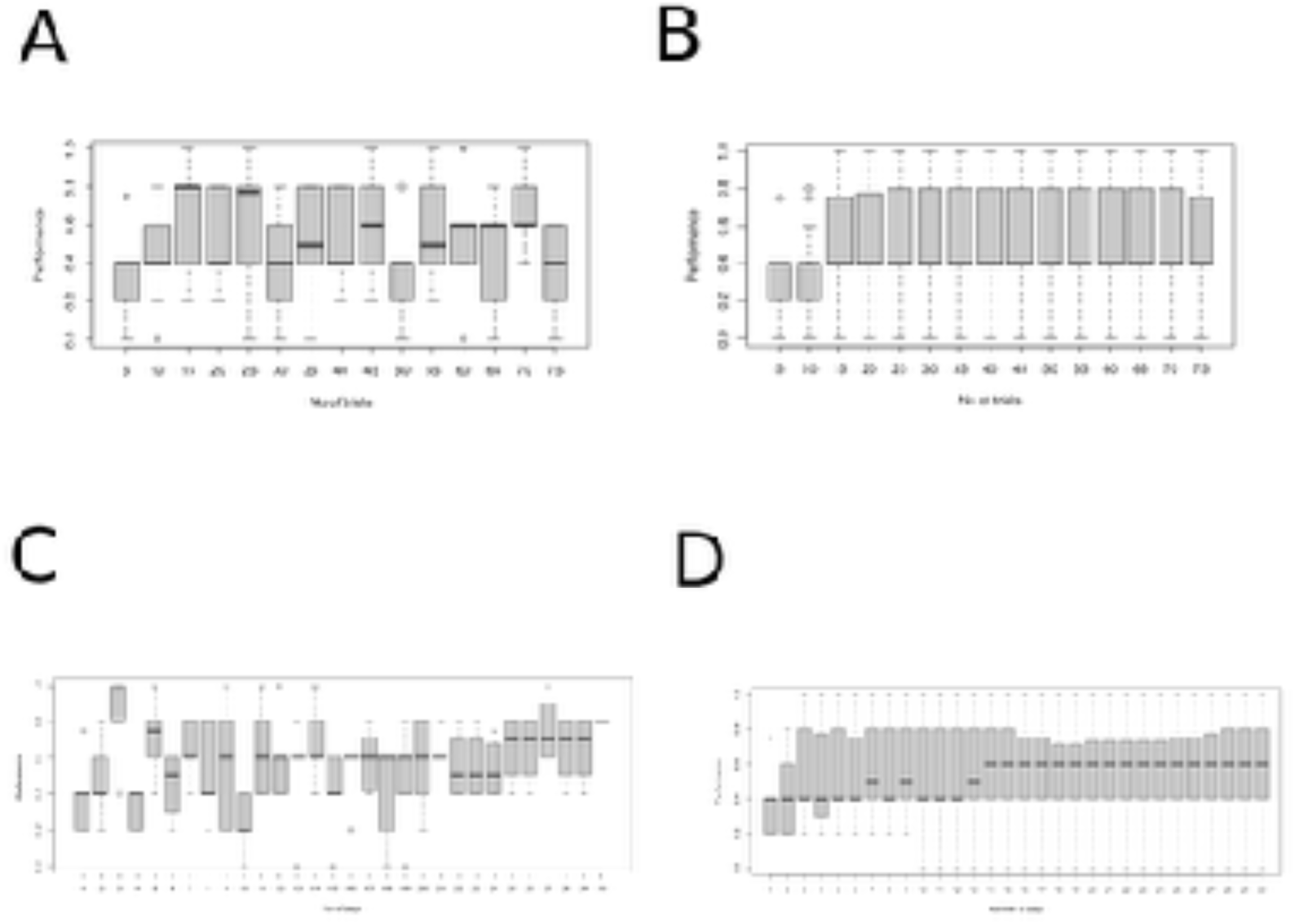
The proportion of trials in which zebrafish choose the side with 2 lines to then receive a reward in the presence/absence of the food reward paradigm across days of training (A) per day and (B) in terms of the cumulative proportion of trials for all fish (n=10). The performance (C) per day and (D) in terms of the cumulative proportion of trials across 21 days of training for a subset of fish who do not show freezing behaviour (n=5).

Since all fish tested in this experiment were individually tracked, we could compare their side bias and WSLS score alongside their choices between the two visual stimuli (data for all five fish shown in Figure 5). Three of these fish reached the learning criterion of statistically significant departure from chance in a binomial test (Figure 5A-C) (66% correct choices over 30 trials, p-value = 0.04937), although a fourth showed an improvement in the proportion of choosing the side with the visual symbol of two lines over the days, that was not statistically significant (Figure 5D). Fish D4 is an example of an individual who did not show any improvement in the proportion of choosing the side with line 2, although it otherwise behaved within the inclusion criteria, with no freezing behaviour or side bias (Figure 5E). These results are interesting because the WSLS spatial learning score is higher than the associative learning performance curve in the individual analysis for all the fish as and when they showed increasing scores for the associative learning task. This suggests that zebrafish may perform well in an associative learning task based on a spatial learning strategy, despite the experimental paradigm involving stimulus side randomization.

**Figure 5:**
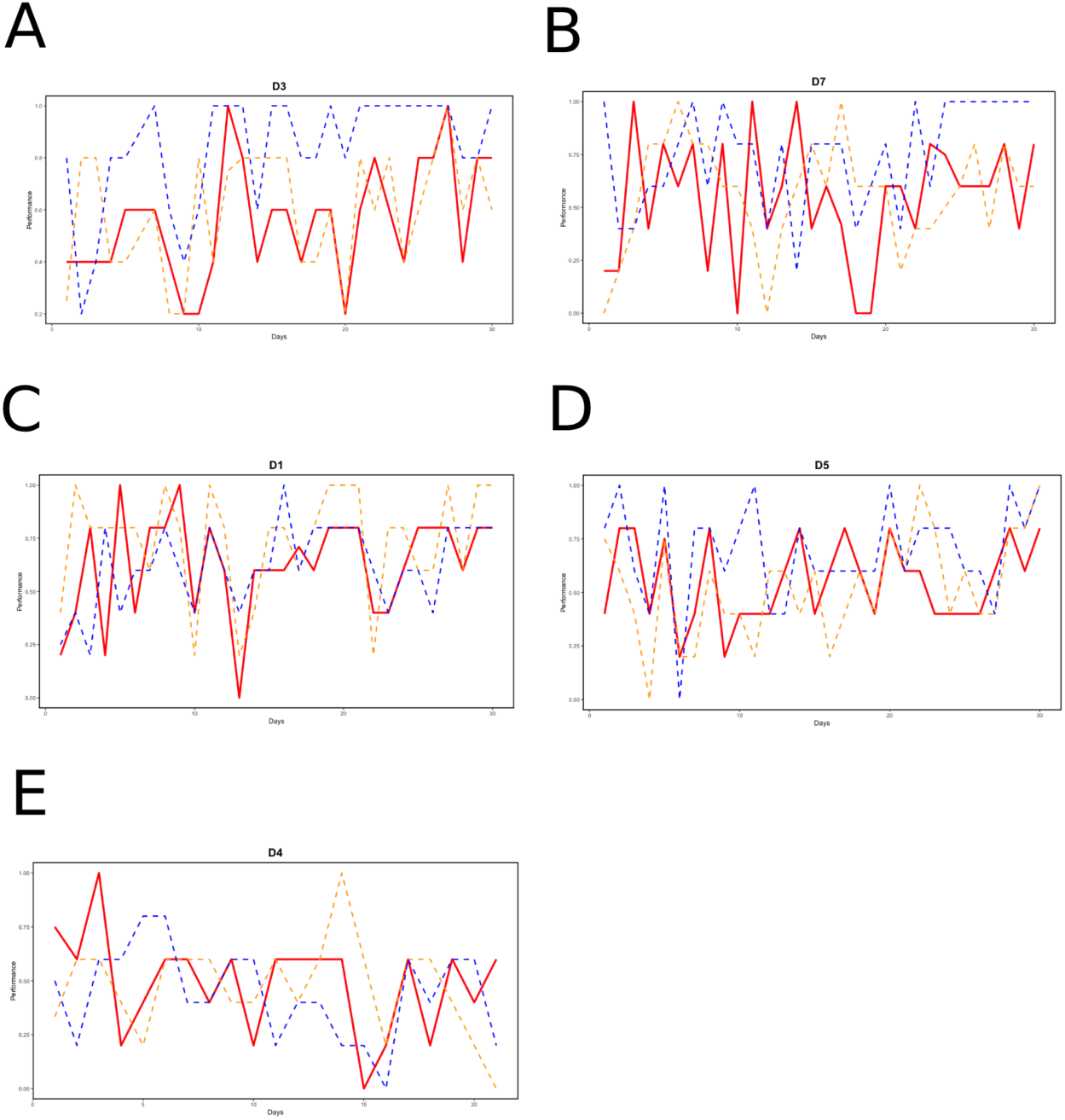
The proportion of trials in which fish choose the side with 2 lines in the presence/ab-sence of food reward paradigm (dark red continuous line), compared with the proportion of trials in which the right side was chosen (orange dotted lines), and the Win-Stay/Lose-Shift scoring metric (blue dotted lines) for individual fish with statistically significant (A, B, C), nonsignificant but improving (D) and worse (E) performance across days of training.

### Experiment 3: Preference for reward quantity in a spatial association task

We performed an additional control experiment to assess food pellet quantity preferences, using a simple spatial association task where the fish were rewarded for going to one side of the T maze with 2 pellets of food, and rewarded when they went to the other side with 1 pellet of food. At the whole population level, zebrafish showed a moderate preference for the larger food pellet quantity (Figure 6A). Two fish showed low interest in food and performed badly in associative learning between spatial location and food pellet quantity, and if these were segregated out of the dataset, a subpopulation of 8 out of 10 fish (Figure 6B) showed individual-level statistically significant learning of this task (all passing the learning criterion of 66% in 30 trials).

**Figure 6:**
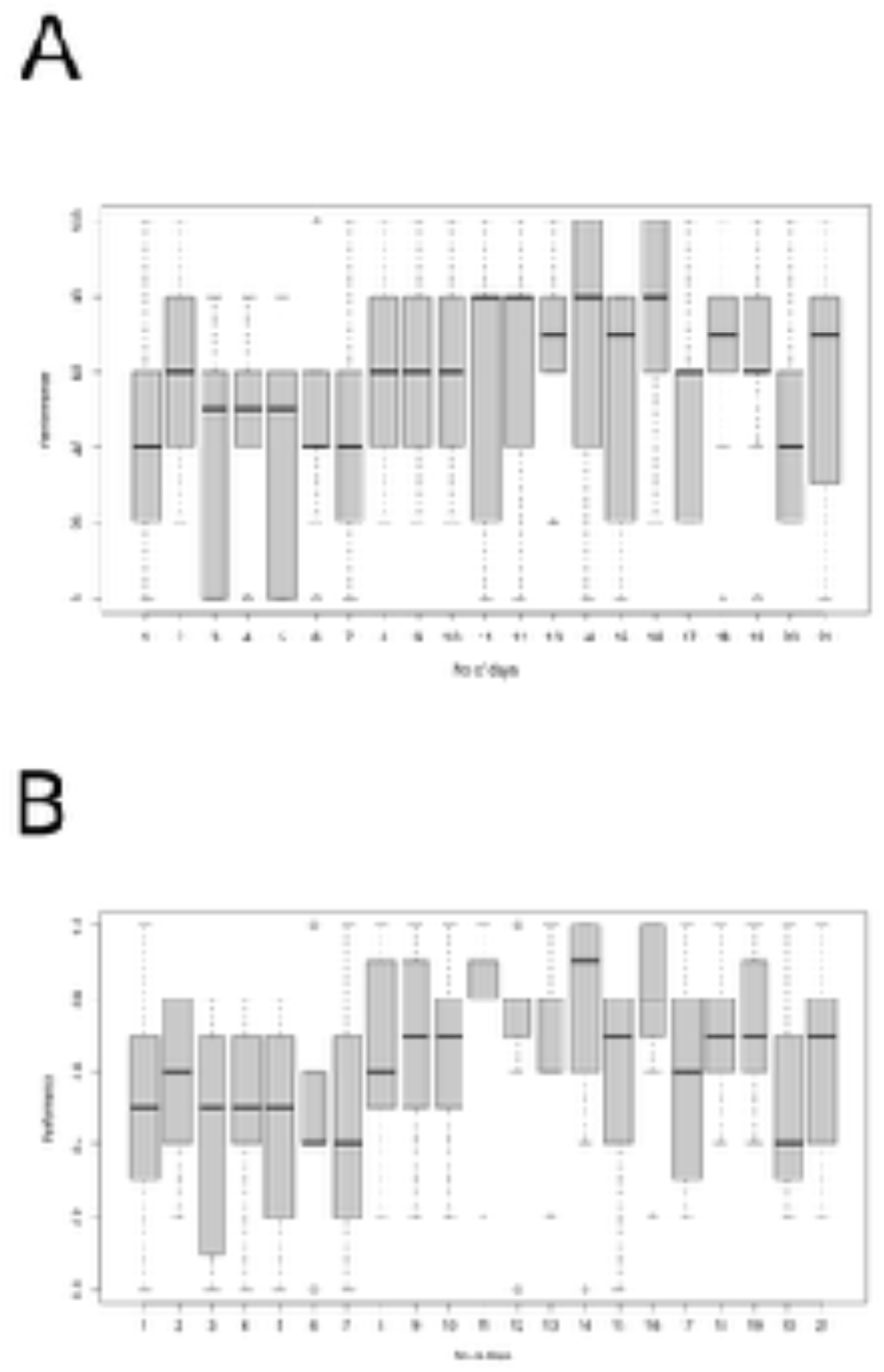
The proportion of trials in which zebrafish choose the side with 2 pellets of food over the side with 1 pellet in a simple spatial association task (A) for all the fish tested (n=10) (B) for a subset of 8 fish all of which individually reached the learning criterion (66% successes in 30 trials).

## Discussion

We carried out three experiments across various levels of task difficulty to investigate associative learning of a visual predictor of reward quantity in wild zebrafish. We found that a majority of the fish were unable to perform the task of learning a predictor of different reward quantities, but that a larger fraction of zebrafish tested could perform a simpler version of the task involving presence/absence of reward. An even larger fraction of zebrafish could learn a further simplified version of the task, without spatial randomization of symbol and reward presentation, such that it was effectively a spatial learning task. Of these, only the latter two tasks have been performed in past zebrafish literature, and therefore they serve as control data establishing the failure to discriminate reward quantity as a feature of wild zebrafish rather than something specific to our setup.

The failure to discriminate between reward quantities must be seen alongside the mixed zebrafish literature reports showing differing results on the quantitative cognitive abilities of the species. The first assessment of the ability of adult zebrafish to associatively learn to discriminate between groups of abstract visual shape stimuli of different cardinalities was part of a comparative study across teleost fish species (Agrillo, Petrazzini, et al., 2012). This inter-species work indicated that far fewer zebrafish reached the learning criteria than the other species tested (guppies, mosquitofish, angelfish, redtail splitfin), and that zebrafish also performed worse than redtail splitfin at a control task involving associating geometric shape discrimination with a reward (Agrillo, Petrazzini, et al., 2012). This study, along with Gatto et al.(2020) suggested that the problem with zebrafish maybe not be with quantitative cognition in particular but with visual shape discrimination for certain kinds of visual stimuli. If so, this would impact performance in any quantitative learning task involving sets of visual stimuli, including our own. However, a recent set of work contests both these findings, suggesting that zebrafish perform well (Bisazza & Santacà, 2022), and equivalently to other fish (guppies (Gatto et al., 2021)) in numerical tasks as well as in tasks involving visual shape discrimination (Santacà et al., 2020).

While food reward discrimination has not been previously assessed, one can argue that spontaneous choices by zebrafish between different group sizes of conspecific fish for sexual assortment constitute an example of quantitative reward discrimination (Potrich et al., 2015; Seguin & Gerlai, 2017). These studies did not control for continuous magnitude information (Potrich et al., 2015) and olfactory information (Seguin & Gerlai, 2017) respectively, and neither did ours. In fact, multiple fish species, including zebrafish, have been shown to spontaneously use continuous magnitude information in shoal size choice (Xiong et al., 2018). If we had found a strong successful symbolic association in this task, our next step would have involved the use of more abstract symbols and reversing the association between these two symbols and the quantity of food they predict, to check whether abstract symbolic visual stimuli can be perceived as predictors of reward magnitude, rather than our non-abstract symbols which bear numerosity and more straightforwardly indicate reward magnitude.

Methodological differences may explain differing outcomes in numerosity experiments on a given model system (Gatto et al., 2017; Howard et al., 2019c), including the salience of the visual stimuli used (Gatto et al., 2020): Bisazza and Santaca (2022) used simpler and more ecologically relevant circular visual stimuli than Agrillo, Petrazzini, et al., (2012), who used a variety of abstract shapes. Associative learning tasks involving more abstract symbols (for example, Arabic numerals or colours/shapes (Howard et al., 2019b; Karoubi et al., 2017)) might be expected to be cognitively more challenging. It was this consideration that prompted us to use simple lines of the kind seen in Roman numerals rather than the more abstract shape stimuli used in tests of cardinality in zebrafish (Agrillo, Petrazzini, et al., 2012; Bisazza & Santacà, 2022) or for similar tests of food reward representation by abstract symbolic shapes in the archerfish (Karoubi et al., 2017). We used lines rather than circles to ensure the fish could see both stimuli while making a choice at the stem of the T maze.

We addressed the question of our results being specific to our setup by testing the same fish on the simpler task of learning the association of the same symbolic stimuli with the presence or absence of the food reward. We found that the zebrafish performed consistently with the findings of Agrillo, Petrazzini, et al., (2012), with less than half of our fish reaching the learning criteria. These inter-individual differences are quite different from the low inter-individual variability and the much higher performance reported by Bisazza and Santaca, with all fish reaching learning criteria (Bisazza & Santacà, 2022). Our use of a T maze and a 2 alternative forced choice paradigm may also be more cognitively challenging compared to the single tank setup used in both of these studies on zebrafish cardinality (Agrillo, Petrazzini, et al., 2012; Bisazza & Santacà, 2022), but we employed a T-maze because zebrafish have been successfully trained to learn to associate a food reward with colour-based cues in a T-maze (Colwill et al., 2005) and with visual and spatial cues in a plus-shaped maze (Sison & Gerlai, 2010). The high performance of fish in our spatial learning task discriminating between 1 and 2 pellets of food confirms that the fish are indeed motivated by this difference in food quantity. It also suggests that spatial learning tasks implemented in a T maze might be cognitively less challenging than a spatially randomized task.

One of the inter-individual differences that emerged related to the degree to which fish were biased towards one side of the T maze, and this is why we explored a spatial learning hypothesis of ‘Win-Stay/Lose-Shift’, where fish make the decision based on the association of spatial cues with the latest positive reward. The criticism (Leibovich et al., 2017b) of the strategy of inferring the perception of numerosity from experiments that sequentially randomize various parameters correlated with numerosity across trials, keeping numerosity constant, also applies to inferring associative learning of stimulus type from tasks where the spatial presentation of the stimulus is randomised across trials. In both cases, sequential randomisation does not preclude the possibility that the animal uses a complex strategy (for example, WSLS in our case) related to the sequentially controlled parameter (in our case, spatial location).

The considerable inter-individual variability documented between individual zebrafish in terms of reaching the learning criteria in associative learning tasks in Agrillo, Petrazzini, et al., (2012) but not in Bisazza and Santaca (2022), was also found in our experiments as well as in several other fish behaviour studies (Budaev et al., 2015; Jones et al., 2018; Karoubi et al., 2017; Lucon-Xiccato & Bisazza, 2017; Lucon-Xiccato & Dadda, 2017; Miletto Petrazzini et al., 2016; Mittelbach et al., 2014; Santacà et al., 2020). Despite reports of inter-individual differences, most work on quantitative discrimination in fish report population-level results and do not provide either individual results, and have not explored spatial learning hypotheses (Agrillo et al., 2009; Agrillo, Petrazzini, et al., 2012; Miletto Petrazzini & Agrillo, 2015; Potrich et al., 2015; Santacà et al., 2020; Xiong et al., 2019). Our study and others in fish (Bisazza & Santacà, 2022; Budaev et al., 2015; Karoubi et al., 2017) and other model systems (Kirschhock et al., 2021; Plotnik et al., 2019; Ramirez-Cardenas et al., 2016) all indicate that individual data could provide valuable insights which could be missed if studies on quantitative tasks in fish ignore heterogeneity within the responses of a population.

Another probable reason for all this variability in zebrafish performance and zebrafish population heterogeneity across labs is likely to be the strains of zebrafish used - all investigations have used local suppliers, but only our suppliers used wild-caught fish from their native habitat in Indian rivers (Arunachalam et al., 2013; Sundin et al., 2019). A high degree of variation in learning and memory behaviours has been demonstrated across different wild-type populations of zebrafish (Roy & Bhat, 2018). It is important to bear in mind while dealing with differences in results across labs (Bisazza & Santacà, 2022, versus Agrillo, Petrazzini, et al., 2012) that different labs working on this question may be working with strains derived from different wild populations, or with strains whose cognitive abilities have diverged considerably from the wild populations they were sourced from (Suurväli et al., 2020; Whiteley et al., 2011). The other possible factor playing into these inter-strain and inter-individual differences might be varying ecologically specific evolutionary advantages of quantitative cognitive abilities. Either way, studies involving wild populations may be more conclusive about the natural range of cognitive abilities of a species as a whole, while those using more derived strains may be more informative about the degree to which high performance may be extracted from a subset of the species.

Our fish were trained for a longer time than most zebrafish associated learning tasks reported in the literature (Agrillo et al., 2009; Agrillo, Petrazzini, et al., 2012; Roy et al., 2019; Roy & Bhat, 2018; Sison & Gerlai, 2010). Some studies stop training when the fish perform significantly higher than the chance level, but we followed a more stringent criterion of waiting for that performance to last for multiple consecutive days due to the inter-day variation in performance (as per Bisazza & Santacà, 2022; Karoubi et al., 2017; Potrich et al., 2019). Even though zebrafish in our second experiment showed significant learning from day 14, we extended the experiment and found that over time, the median performance in doing the task which had reached 0.6 (the performance documented in Agrillo, Petrazzini, et al., 2012)), increased further to 0.7 (as documented in Bisazza & Santacà, 2022) along with reduced variability. Prolonged training can therefore elevate the performance in associative learning tasks in zebrafish. The question of whether prolonged training can elevate performance was also raised by Bisazza and Santaca (2022) in comparing the relatively low accuracy in results of training fish, with experiments involving thousands of trials in birds and mammals where the accuracy is closer to 90-95%; and in fact, such extensive training has been shown to raise the performance of goldfish beyond 90% with 1200 trials (Bisazza et al., 2014; DeLong et al., 2017).

Zebrafish might be able to perform well on a quantitative symbolic learning task that is structured differently. While archerfish can perform very well in similar tasks associating a visual stimulus with reward quantity (Karoubi et al., 2017), work on bees suggests they can learn to associate abstract symbols representing number with visual stimuli consisting of a set of the same number of objects, for a reward/no reward (Howard et al., 2019b). We found that a larger fraction of zebrafish was able to associate each of two symbolic visual stimuli with the presence/absence of a food reward respectively, than with different quantities of food. An even larger fraction was able to perform statistically significantly above chance levels at a spatial learning task that involved discriminating between the same quantities of food. This suggests different degrees of ease of performance and high inter-individual variation in the ability of wild zebrafish to perform the harder cognitive tasks to which we subjected them. Potrich et al., (2024) tested a similar gradation of task difficulty with respect to arithmetic ability and the preference of zebrafish for larger social groups, and found zebrafish only showed preferences for the larger group in the #1 vs. 2” comparison, but not in the more demanding #2 vs. 3” or #2 vs. 4” conditions,.

This investigation of symbolic quantitative cognition is a necessary step to examine the ability of zebrafish to perform associative quantitative operations beyond the presence/absence-of-reward paradigm that has largely been used in associative learning tasks with quantitative symbols. This broadens our understanding of the cognitive abilities of this model system in which neuroimaging can cast a wide net across the brain in examining the neural and genetic basis of associative quantitative learning abilities and clinical variations such as dyscalculia (Bisazza & Santacà, 2022; Messina et al., 2022).

This work could be taken forward by examining symbolic quantitative cognition in zebrafish using other testing paradigms involving social rewards, which can sometimes be more potent than food rewards (Karnik & Gerlai, 2012; Lucon-Xiccato et al., 2015). Given the general performance of zebrafish on visual shape discrimination, symbolic quantitative cognition could be tested using other modalities such as auditory stimuli. It is also clear that all quantitative learning results that use spatial randomization must be considered in the light of alternate hypotheses such as spatial side bias and spatial win-stay lose-shift strategies: further experimentation would be necessary to examine zebrafish associative learning tasks to distinguish between these two hypotheses explaining their results. We have also used wild-type zebrafish caught from their natural habitats in India and it would be valuable to conduct a comparative study looking at the population and/or strain specificity in these quantitative cognitive abilities such as has been conducted on learning and memory (Roy & Bhat, 2018). Finally, work on this model system raises the possibility of using neuroimaging and genetic techniques to understand the neural and genetic basis of non-symbolic and symbolic quantitative cognition abilities and disabilities.

## Acknowledgements

This research was supported by the Ashoka University Annual Faculty Research grant and the infrastructure provided by the Research and Development Office, and the DST-INSPIRE grant. The authors would like to thank L.S. Shashidhara, Krishna Melnattur, Joby Joseph for discussions and feedback on the experiments and the manuscript.

